# Autoregulation of three yeast ribosomal protein genes by splicing inhibition

**DOI:** 10.64898/2025.12.19.695493

**Authors:** David Granas, Anirudh Kesanapally, Michael A. White, Gary D. Stormo

**Affiliations:** Department of Genetics and Center for Genomics and Systems Biology, Washington University School of Medicine

**Keywords:** autoregulation, mRNA structure, splicing

## Abstract

Yeast ribosomal protein genes *RPL18B, RPL28* and *RPS22B* are autoregulated by inhibition of splicing. This is demonstrated by inserting their introns into a chromosomal copy of GFP and observing repression upon induction of the cognate protein. In *RPL18B* and *RPS22B*, a predicted conserved secondary structure within the intron is required for regulation, while in *RPL28* it is not.

## INTRODUCTION

Autoregulation of gene expression is an efficient means of maintaining protein concentrations in a narrow range (Savageau 1974; Alon 2007; Crews and Pearson 2009). It is common among transcription factors (TFs) because it is readily evolved through the acquisition of a binding site for the TF within the regulatory region controlling the gene’s expression (Bateman 1998). Those non-coding regions can evolve rapidly and if the autoregulation confers an advantage, which is often the case, it can be selected for. Post-transcriptional autoregulation is also common for RNA-binding proteins where acquisition of binding sites in regions that control translation, or splicing or mRNA degradation, can also be selected for (Yeo *et al*. 2007; Buratti and Baralle 2011; Muller-McNicoll *et al*. 2019). It has even been observed for proteins whose normal function does not include regulation of gene expression, where the only affected gene is that encoding the protein (Russel *et al*. 1976; Andrake *et al*. 1988; Torres-Larios *et al*. 2002; Romby and Springer 2003).

In previous work we used the yeast GFP collection (Ghaemmaghami *et al*. 2003) to identify many yeast ribosomal protein genes that showed feedback regulation of expression (Roy *et al*. 2020). We showed that the mechanism of autoregulation for *RPL1B* was by inhibiting translation initiation through binding to a conserved sequence and structure motif just 5’ to the initiation codon. That mechanism is similar to many examples in bacteria (Nomura *et al*. 1980; Nomura *et al*. 1984; Draper 1989; Deiorio-Haggar *et al*. 2013). We next wanted to identify new examples of autoregulation that involved inhibition of splicing, a mechanism not available in bacteria. We used four criteria to pick genes to test. First, they had to have an intron near the 5’ end of the gene, as that is expected to be the most likely regulatory target. And second, the intron had to contain a predicted secondary structure that is conserved across multiple yeast species (Hooks *et al*. 2016). Third, for genes included in our GFP collection (*RPL28* was not) they had to show feedback regulation in our previous study of GFP-tagged proteins (Roy *et al*. 2020). Fourth, in the study by (Gelperin *et al*. 2005), whole genes, including introns, were cloned for high expression under the control of the *GAL1* promoter. Genes with low or moderate expression are candidates for autoregulation. This search resulted in four genes that we suspected might be autoregulated by inhibition of splicing: *RPS22B, RPL7B, RPL18B*, and *RPL28. RPL7B* was analyzed in detail and is described elsewhere (Granas *et al*. 2025). In this manuscript we describe our findings for the other three genes. We reasoned that if the intron contains the cis-regulatory site, then placing the intron in the GFP gene will repress GFP expression following over-expression of the cognate protein.

## MATERIALS and METHODS

### Yeast strains

Yeast strains were made with a chromosomally-integrated copy of eGFP containing different introns. The wild-type intron sequences were synthesized by either IDT (*RPS22B*) or Synbio (*RPL18B, RPL28*). Upstream sequence before the intron for each gene was also included (5’ UTR and CDS for *RPL18B* and *RPL28*, and 5’ UTR for *RPS22B*). These constructs were cloned into the plasmid BAC690-TEF (described in Roy *et al*. 2020) using NEB’s HiFi DNA Assembly. Intron variants were then made using NEB’s Site-Directed Mutagenesis Kit. A PCR amplicon was made from this plasmid containing the *TEF2* promoter, eGFP with the upstream sequence and intron, the ADH1 terminator, and the kanamycin/G418 resistance marker, all flanked by homology arms to target integration to the dubious ORF YBR032W. This PCR amplicon was transformed into the FM391 yeast strain (genotype: MATa, his3Δ leu2Δ ura3Δ met15Δ) using the LiAc/SS carrier DNA/PEG method. Transformants were selected by growth on YPD plates containing 400 μg/mL G418 and then verified by sequencing.

Plasmids MJB1-Rpl18, MJB1-Rpl28, and MJB1-Rps22 were synthesized by Twist Bioscience to express the cre-less genes under control of the inducible *GAL1* promoter. For Rps22 versions were made with both an N-terminal mCherry tag and no tag, while Rpl18 and Rpl28 both used C-terminal mCherry tags. These plasmids contain a CEN/ARS element and the *URA3* gene for selection. The plasmids were then transformed into the strains containing the genome-integrated eGFP/intron construct and grown on SC-URA plates. The sequences of the cre-less genes and proteins are provided in Supplemental Table T1.

### GFP assays

Yeast colonies were inoculated into 400 μL SC-URA media with 2% raffinose and grown overnight at 30 °C. The next day cultures were diluted into both SC-URA with 2% raffinose (uninduced) and SC-URA with 2% raffinose and 0.2% galactose (induced). Cells were grown at 30 °C for 10 hours and then assayed on a CytoFLEX S (Beckman Coulter). Live cells were gated and 10,000 events were acquired. GFP was measured at an excitation of 488 nm with a 525/40 filter.

## RESULTS and DISCUSSION

Our approach is described in Figure 1. A “cre-less” version of gene X (each of the genes being studied) is placed under control of the *GAL1* promoter and is therefore inducible by the addition of galactose to the media. “Cre-less” refers to the fact that all potential cis-regulatory elements of the gene have been removed: the 5’ and 3’ UTRs have been replaced, the introns have been removed and the codons have been shuffled to code for the same protein using a different DNA sequence (Roy *et al*. 2020). GFP has been inserted into the genome under control of the constitutive *TEF2* promoter (Peng *et al*. 2015). The intron, or variants of the intron, for gene X are inserted 5’ of GFP so that splicing is required for GFP expression. Sequences upstream of the intron (the 5’ UTR and coding sequences) are also included. If GFP expression is reduced upon induction, that shows that the insert is a cis-regulatory site under the control of the protein for gene X. The simplest explanation is that protein X binds to the intron to inhibit splicing, although more complex, perhaps indirect, mechanisms cannot be ruled out.

**Figure 1.**
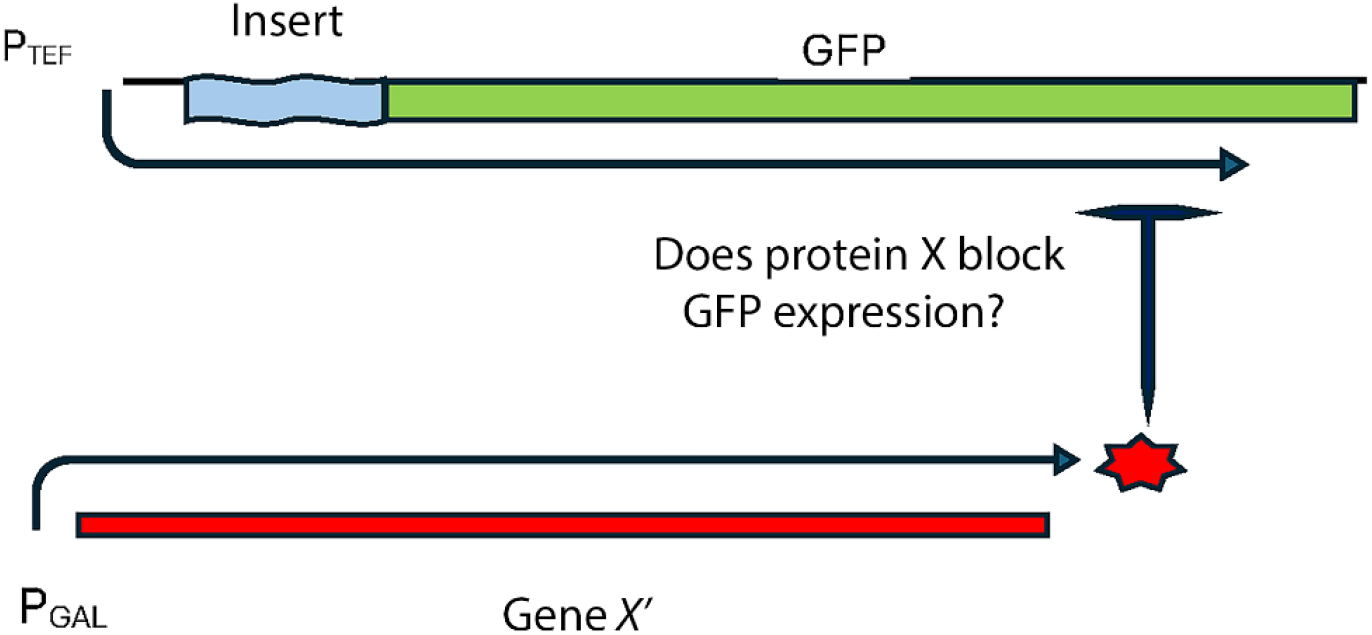
Strategy for measuring regulatory effects. A “cre-less” version of each gene (*RPL18A, RPL28, RPS22A*), referred to as X’, was cloned into a plasmid where it could be induced to express the protein X. The “insert” is the intron for gene X and various mutated versions of it that are placed 5’ of the chromosomal GFP driven by the constitutive TEF promoter. Upstream sequences of the gene (including 5’ UTR and 5’ CDS) were also included in the insert. GFP levels are measured with and without induction to determine if each insert contains a cis-regulatory site acted upon by protein X.

Figure 2 a,b,c show the change in fluorescence for each of the wild-type introns after induction of the cognate gene. Table 1 lists the change in expression in column 2 for each gene listed in column 1. In every case expression is reduced indicating the intron is a site for autoregulation, varying from about 12.5 fold (log value 1.1) for the first intron of *RPS22B*, to about 3 fold (log 0.5) for *RPL18B*. Those results show that we successfully predicted that the intron contains the cis-regulatory site for autoregulation of each gene. Figure 2 d,e,f show the change in expression after induction if the Hooks structure has been deleted from the intron, and the change values are listed in column 3 of Table 1. By Hooks structure we mean the secondary structures identified by (Hooks *et al*. 2016) using conservation-based computational predictions. For *RPS22B* and *RPL18B* the inhibition of expression is almost completely eliminated, indicating that the Hooks structure is required for autoregulation. For *RPL28* regulation is maintained after deletion of the Hooks structure. The results for each gene are described in more detail below. GFP assays for all of the intron variants, and their sequences, are provided in Supplemental Table S2.

**Table 1.** Log change in expression after induction of the cre-less gene. For the full intron and with a deletion of the Hooks structure.

| Intron | Full log(unind/ind) | Del-str log(unind/ind) |
| --- | --- | --- |
| Rps22b* | 1.1 | 0.1 |
| Rpl18b | 0.5 | 0 |
| Rpl28 | 0.8 | 0.7 |

**Figure 2.**
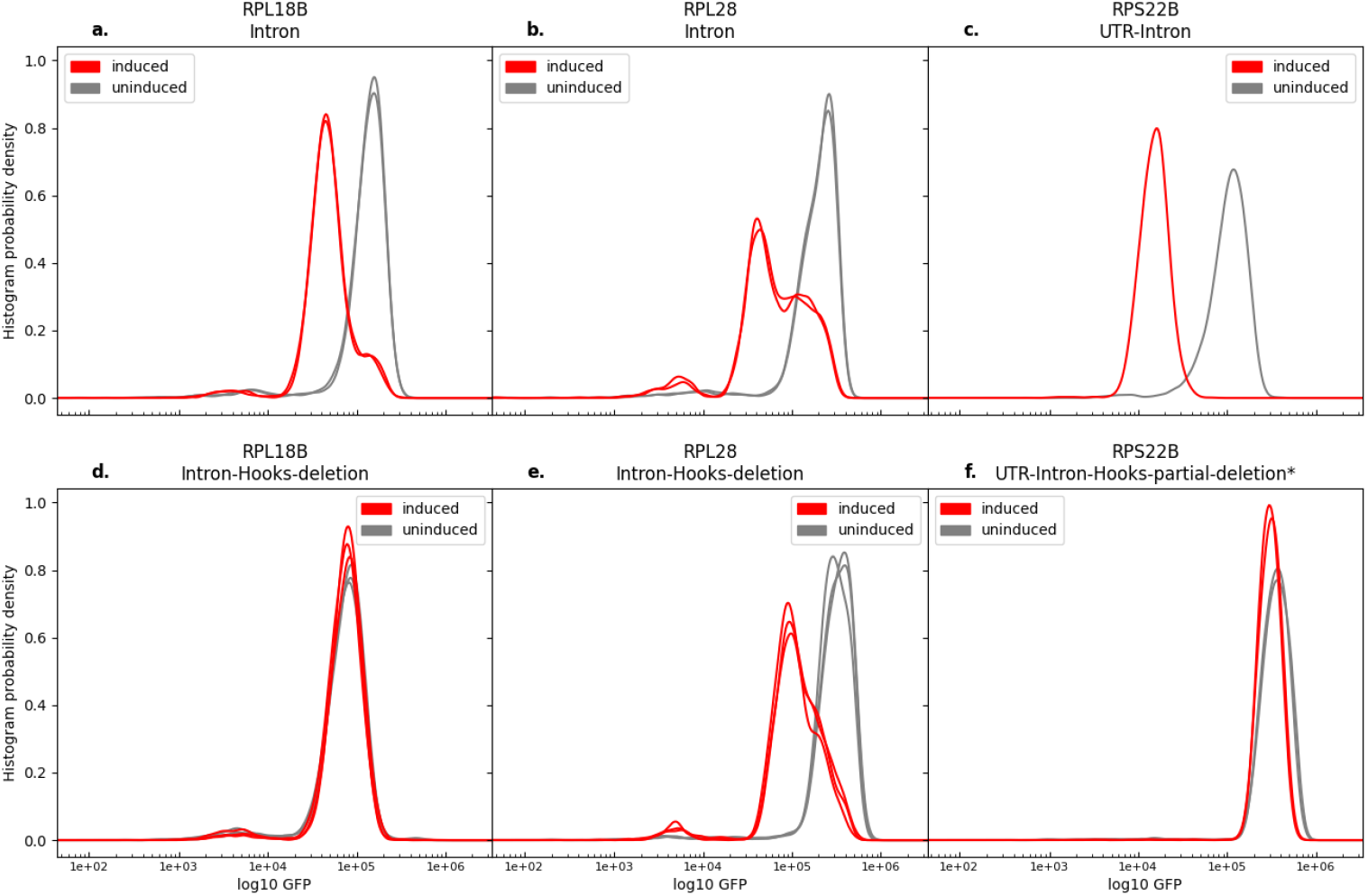
GFP measurements with and without induction. a.-c. The inserts contain the full introns of the specified gene. d.-f. The inserted introns are deleted for the Hooks structure. For *RPS22B*, only a portion of the Hooks structure was deleted, that shown as S1 in Figure 3b.

### RPL18B

*RPL18B* and *RPL18A* have a conserved secondary structure (Hooks *et al*. 2016). *RPL18A* has an additional structure, not conserved in *RPL18B*, that serves as an RNaseIII (Rnt1p) cut site (Danin-Kreiselman *et al*. 2003). Both introns show regulation upon induction of the Rpl18 protein, but expression is only reduced about 20% for *RPL18A* (not shown). For *RPL18B* expression is reduced about 3-fold (Figure 2a, Table 1). Deletion of the Hooks structure in *RPL18B* reduces expression by about one-third from the wild-type level, and it is no longer regulated by overexpression of the Rpl18 protein (Figure 2d, Table 1). In this case the Hooks structure is required for regulation but also contributes to the normal, unregulated, level of expression. We did not pursue this intron further.

### RPL28

*RPL28* in *S. cerevisiae* is the homolog of L15 in bacteria and is also known as uL15 (Ban *et al*. 2014). The gene is 961 bases long from start codon to stop codon, with a 511 base long intron beginning at position 50 after the start codon. That intron contains the predicted Hooks secondary structure of 116 bases beginning at position 119 from the 5’ splice site (Hooks *et al*. 2016). The branch point sequence, TACTAAC, starts 228 bases after the Hooks structure and ends 42 bases before the 3’ splice site. As shown in Figure 2b and Table 1, upon induction of the Rpl28 protein, GFP expression is reduced about 6-fold, although there is a distinct shoulder in the graph, indicating not all cells show strong inhibition. As shown in Figure 2e, when the Hooks structure is deleted, repression still takes place, and now the shoulder is much reduced. These results indicate that the intron contains a cis-regulatory site that represses expression when the cognate protein is over-expressed, presumably by inhibiting splicing, but that the Hooks structure is not required for the regulation. Rangan et al (Rangan *et al*. 2025) have analyzed yeast intron structures using a combination of *in vivo* structure probing and computational analysis. Their model for the *RPL28* intron contains a structure very similar to, and overlapping, the Hooks structure, that is located at the end of a long stem. In a model of the intron interacting with the spliceosome, that region of the intron is far removed from the spliceosomal complex, consistent with its deletion not affecting expression or regulation.

The initial insert for *RPL28* contains 16 bases of the 5’ UTR, the first 49 bases of the coding sequence, the intron, and 8 more bases of the coding sequence (Supplemental Figure S1, V1). Since the Hooks structure is not required for regulation, we thought that perhaps sequences outside the intron may contribute. Therefore we made additional variants of the intron, shown in Supplemental Figure S1. Supplemental Figure S2 shows the GFP signal before and after induction for each of those insert variants. The results show that the 5’ UTR is not required for regulation, and that neither the intron alone nor the coding region alone is sufficient for regulation. At least some portion of the intron and some portion of the coding region are required for regulation. We suspect that the binding site for Rpl28p may be a sequence and structure motif overlapping the 5’ splice site, as in the case of RPL30 regulation (Vilardell *et al*. 2000), or perhaps the 3’ splice site.

Inducing Rpl28p expression causes a growth defect in cells as evidenced by lower cell density after 10 hours of induction compared to the uninduced cells. This does not depend on the *RPL28* intron, as it also happens when the *RPL7B or RPS22B* introns are upstream of GFP when Rpl28p is induced. In those cases there is no regulation of GFP, but the growth rate is reduced. Measurement of mCherry by flow cytometry shows that there are always some cells that are not induced with the addition of galactose to the media, presumably because they have lost the plasmid or it has picked up mutations that prevent expression of cre-less *RPL28*. When we measure GFP we “gate” on the level of mCherry, so only cells that show induction (an increase in mCherry over background) are included. When *RPL28* is induced, not only do cells grow slower, but a larger fraction of them do not show any induction, indicating a selection against overexpression. There are also cells with mCherry signal that is low, but above background, and there is a correlation between cells with low mCherry expression and those with the GFP shoulder, suggesting that lower Rpl28p levels leads to less regulation. Supplemental Figure S3 shows 2D plots of GFP and mCherry before and after induction for both inserts V1, which shows regulation, and V2, which does not. Using the upper level of mCherry in the uninduced state as the cutoff, we see that after induction there is a wide range of mCherry expression with both inserts. With insert V2 there is no reduction in GFP, whereas with insert V1 some cells show reduced GFP and others do not, corresponding to the regulated peak and the shoulder we observe. We have not pursued this intron further.

### RPS22B

*RPS22B* has two introns, including one entirely within the 5’ UTR. We previously showed that the intron within the 5’ UTR is sufficient to confer repression upon induction of the Rps22 protein (Roy *et al*. 2020), which is confirmed again in Figure 2c and Table 1. That intron contains a conserved predicted structure (Hooks *et al*. 2016) within which is an RNaseIII (Rnt1p) cut site, in approximately the middle of the conserved Hooks structure (Danin-Kreiselman *et al*. 2003). When part of the Hooks structure is deleted, expression increases about 3-fold over the strain with the wild-type intron, but regulation is completely lost (compare Figures 2c and 2f, Table 1). To determine what parts of the intron are required for regulation we made several variants of the intron and measured their expression, shown in Figures 3 and 4.

**Figure 3.**
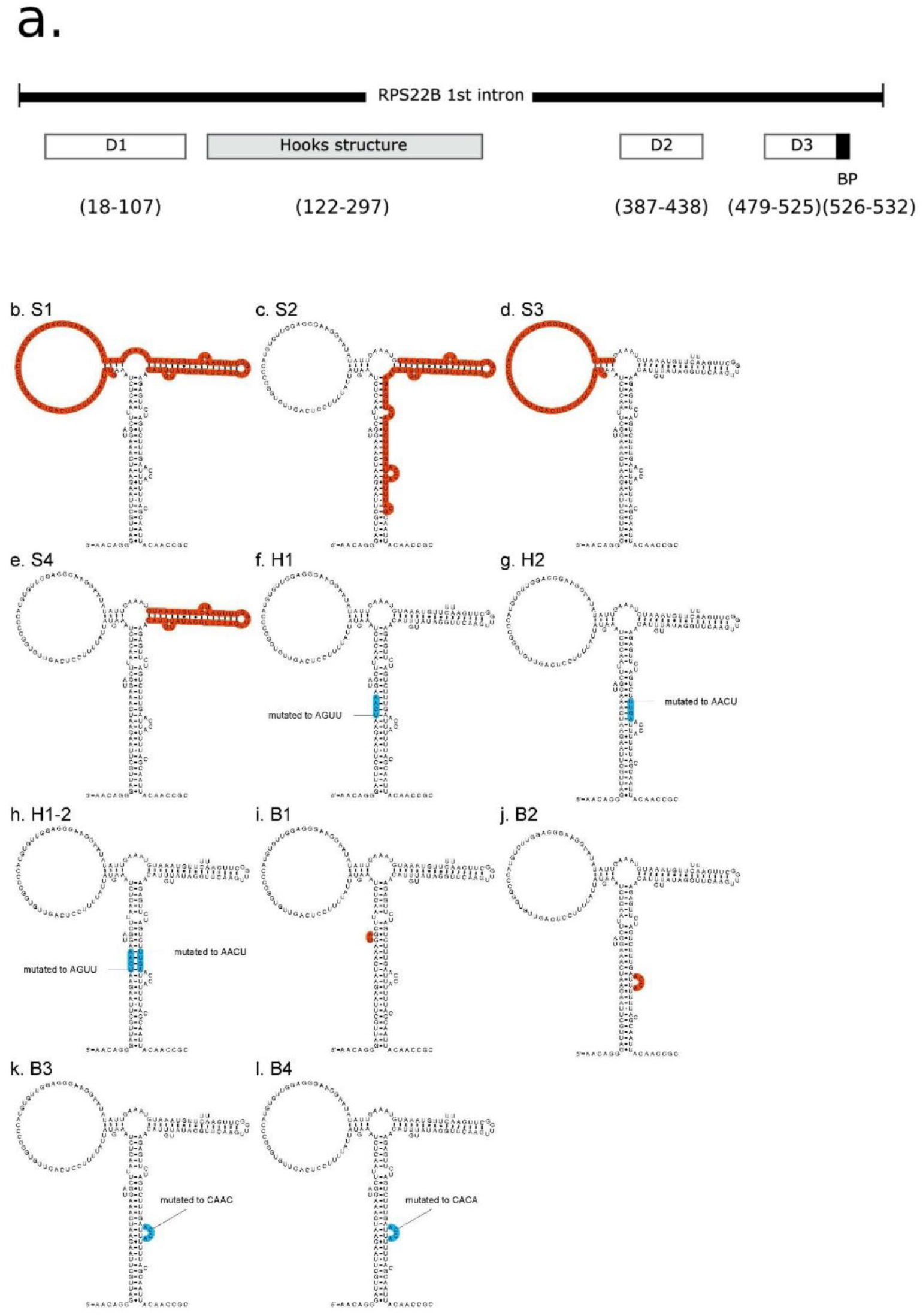
*RPS22B* intron 1 and mutated variants. a. A schematic of the entire intron, showing the positions of deletions D1-D3 and of the Hooks structure. b.-l. Each of the variants of the Hooks structure. Red sections were deleted and blue sections were mutated to the indicated sequences.

**Figure 4.**
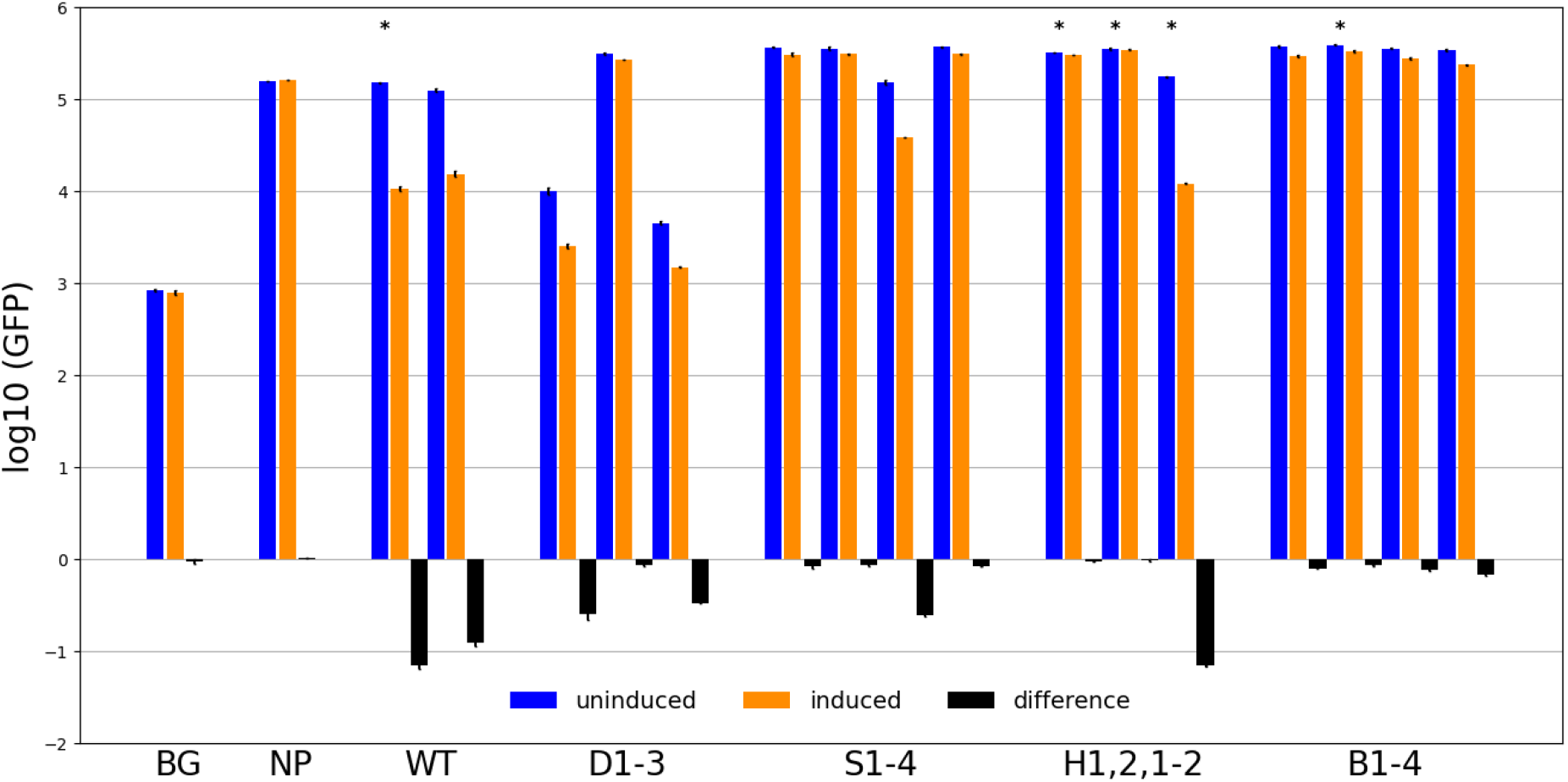
GFP expression values of *RPS22B* intron variants. The vertical axis is the log (GFP fluorescence) with error bars indicating the standard deviations. Blue bars are from cells in the uninduced state, orange bars are for the induced state, and black bars are the difference (induced – uninduced). BG is the background fluorescence in cells without GFP. NP is cells that have the wild-type 5’ UTR intron from *RPS22B* upstream of GFP but do not have Rps22 on the plasmid to be induced (instead only mCherry is induced), showing that the change of media to contain galactose has no effect on GFP fluorescence. WT is cells that have the wild-type 5’ UTR intron of *RPS22B* upstream of GFP and also have the induction plasmid containing the cre-less Rps22 protein. The others correspond to the variants shown in Figure 3b-l. Bars with * above them were done using an untagged version of the Rsp22 protein, whereas the others have mCherry fused to the N-terminal end of the protein.

Figure 3a is a schematic of the *RPS22B* intron from the 5’ UTR. It is 557 bases long with the 180 base Hooks structure occurring 122 bases after the 5’ splice site. We made three deletions, labeled D1-3 in Figure 3a, that are outside of the Hooks structure, D1 on the 5’ side and D2 and D3 on the 3’ side. Figures 3b-l show variants made to the Hooks structure that were tested for repression activity. Regions shown in red were deleted, while those shown in blue were mutated to the indicated sequence. Deletions S1-4 remove large segments of the Hooks structure. H1 and H2 are changes to the sequence of the central stem and in the combination H1-2 the stem is reformed with the alternative sequences. We also deleted the two bulge loops, the small UA on the left (B1) and the larger one of the right (B2), and we changed the sequence of the larger loop in two different ways (B3,B4).

Figure 4 shows the measured log(GFP) for all of the strains, both in normal media and in inducing media that contains galactose. Standard deviations from replicate experiments are shown on each bar and indicate the very high reproducibility of the assays. BG is the background fluorescence in cells without any GFP, showing the inherent fluorescence in the yeast strain we use (FM391). NP is a strain with the wild-type intron in GFP but no *RPS22* gene on the plasmid, showing that changing the media does not affect the fluorescence signal. WT refers to replicate experiments with the wild-type intron upstream of GFP. Bars labeled with (*) do not have an mCherry tag on the Rps22 protein. In our original study we intended to fuse each cre-less gene to mCherry on the C-terminus to monitor their expression. Inadvertently, the stop codon was left on the RPS22 gene, so there was no mCherry tag. That turned out to be fortunate because when the mCherry fusion was made to the C-terminus, Rps22p was inactive (at least for regulation). We then tested if an N-terminal fusion is active, and it is. Most experiments reported here were performed with the N-terminal mCherry, but a few, marked with *, were with the untagged version of the protein. As can be seen in replicates with the wild-type intron, the N-terminal tagged version and the untagged version of the protein behave almost identically, except that the untagged version has a slightly larger degree of repression (log difference = −1.15 vs −0.9). We suspect this is due to more of the untagged protein being available, either because it is translated more efficiently or is degraded less rapidly.

Deletions D1 and D3, near the 5’ end of the intron and the 3’ end on the intron, respectively, reduce expression in the uninduced state by over 10-fold. We suspect there is a “zipper stem” that brings the ends of the intron closer together, as is seen in many other ribosomal gene introns (Rangan *et al*. 2025). Both deletions maintain regulation but at a lower level, about 3-4 fold. Deletion D2, 3’ of and near to the Hooks structure, is expressed about 3-fold higher than the wild-type intron in the uninduced state and has lost regulation. In fact, every intron variant that has lost regulation is expressed about 3-fold higher than wild-type in the uninduced state. We infer that the endogenous level of Rps22p, from the nascent genes, limits expression, and when the regulatory site is removed that regulation is eliminated.

Deletions S1, S2 and S4, of parts of the Hooks structure, all lose regulation and are expressed at the higher level in the uninduced state. Deletion S3, which deletes the large loop on the left of the Hooks structure shown in Figure 3d, is expressed at near wild-type levels in the uninduced state and is still regulated at a lower level of about 4-fold. If the central stem of the Hooks structure is eliminated by mutating either the 5’ or 3’ sequence (H1 and H2), expression is increased and regulation is eliminated. But if both mutations occur (H1-2), which regenerates that central stem with altered sequence, then expression returns to normal as does regulation.

All of the mutations to the two bulge loops in the stem of the Hooks structure increase expression and eliminate regulation. That was the case even of the two mutations that left the large bulge in place but changed the sequence (B3,B4), indicating that both the structure and the sequence are required for function. Rps22 is the homolog of bacterial protein S8 (also known as uS8 (Ban *et al*. 2014)) which, *in E. coli*, regulates the *spc* operon containing 10 ribosomal protein genes (Cerretti *et al*. 1983; Cerretti *et al*. 1988; Mattheakis and Nomura 1988). The X-ray crystal structure of the *E. coli* S8 protein bound to the regulatory site on the mRNA, which overlaps the 5’ end of the L5 gene on the polycistronic mRNA, has been determined and compared to the crystal structures of S8 bound to the small subunit rRNA in ribosomes from several bacteria (Merianos *et al*. 2004) and to an NMR structure of S8 bound to helix 21 of 16S rRNA where it binds (Kalurachchi *et al*. 1997). The RNA sequences of those differ but they all contain a stem with two bulge loops, similar to the predicted Hooks structure of *RPS22B* shown in Figure 3. However, the actual 3-D structures of those RNA segments are more complicated than the simple 2-D base-pairing diagram. Bases from the two bulge loops fold back to interact with base-pairs in the stem to form two base-triple interactions stacked atop each other. We suspect the central region of the Hooks stem in *Rps22B* can form a similar complex structure with the two bulge loops, but we have not been able to infer an alignment between them. We showed that disruptions of the central stem eliminate regulation, as do deletions of either bulge loop or changes to the sequence of the larger one. Inferring complex 3-D structures of RNA from sequence is a challenging problem and thus we cannot say whether the Hooks structure resembles the structure bound by S8 in bacteria.

If the central stem constitutes the binding site for Rps22, it is not clear why deletions S1 or S4 should eliminate regulation. S3 is still regulated, to a lower extent, which is consistent with the model of the central stem being the primary binding site. One possibility is that those deletions alter the structure of the intron so that the Rps22p binding site is no longer available. Another possibility is that the Rnt1p (yeast ortholog of RNaseIII) plays a role in regulation. It was shown that the stem deleted in S4 is the cut site for Rnt1p (Danin-Kreiselman *et al*. 2003), and in strains deleted for *Rnt1*, or with deletions to the intron equivalent to our S1 and S4, *RPS22B* mRNA levels were increased about 4-fold, similar to our observation for all of the non-regulated variants. They hypothesized that *Rnt1* had two roles, one was to initiate degradation of the intron lariat after splicing, and the other was to cleave non-productive, unspliced pre-mRNA. But it is intriguing that deletion S2, which removes both the central stem and the Rnt1p cut site, has exactly the same phenotype as removing either site alone. An interesting possibility is that Rps22p and Rnt1p function cooperatively to control the fate of the intron. In this model, when Rps22p is in excess, it binds to the *RPS22B* intron along with Rnt1p to initiate cleavage and degradation of the intron, thereby autoregulating its expression. Even if that model is correct, it does not explain the role for the sequence deleted in D2, and why that is required for regulation. One possibility is that the Rps22 protein interacts with a larger segment of RNA than just that central stem, which would be different from how the bacterial S8 proteins interact with RNA. Another possibility is that deletions of segments of RNA can have large effects on RNA structures outside of where the segment is located. Unlike DNA, where deletion of a segment of sequence can be reasonably assumed to only affect the sequence of the DNA and not the structure, deletions of any RNA segment, or even just mutations of any bases, may lead to significant changes in the overall structure. Therefore, while we suspect that Rps22 protein interacts with the intronic RNA in a manner very similar to how bacterial S8 protein interacts with its RNA binding sequences, we cannot rule out more complicated models. Additional effort will be required to answer that question.

## CONCLUSIONS

Most yeast ribosomal genes (>70%) contain introns, in contrast to other genes where introns are rare (<5%). In addition, ribosomal gene introns are substantially longer than the other introns (Spingola *et al*. 1999). Rangan et al (Rangan *et al*. 2025) have examined yeast intron structures using *in vivo* structure probing experiments and computational modeling. They find that most ribosomal gene introns contain “zipper stems” that bring the 5’ end of the intron near to the branch point sequence, effectively making the intron shorter and presumably more efficiently spliced. In fact, previous work has shown that intron structures can serve as enhancers for splicing (Libri *et al*. 1995; Granas *et al*. 2025).

An extensive analysis of ribosomal genes showed that many intron deletions had growth phenotypes under some conditions (Parenteau *et al*. 2011). Our examples add to the list of introns that have a role in controlling gene expression, specifically through autoregulation of splicing. There are likely many more examples. Of the 95 ribosomal protein genes in our GFP collection, 30 of them had at least a 3-fold reduction in GFP expression when the cognate protein was over-expressed, nearly all of which were novel findings (Roy *et al*. 2020). Of course, that approach does not demonstrate the mechanism of feedback regulation. It could be transcriptional, or effect any of many post-transcriptional steps leading to protein production, or even enhanced protein degradation of the excess proteins. Given the potential for RNAs to be sensors of the cellular environment, we expect there to be many more examples of their role in regulating gene expression, not only autoregulation but cross-regulation as well.

## Data availability

The flow cytometry measurements are available at https://figshare.com/articles/dataset/RPL18B_RPL28_RPS22B_splicing_autoregulation/30916613

