## supplemental figures for "Autoregulation of three yeast ribosomal protein genes by splicing inhibition"

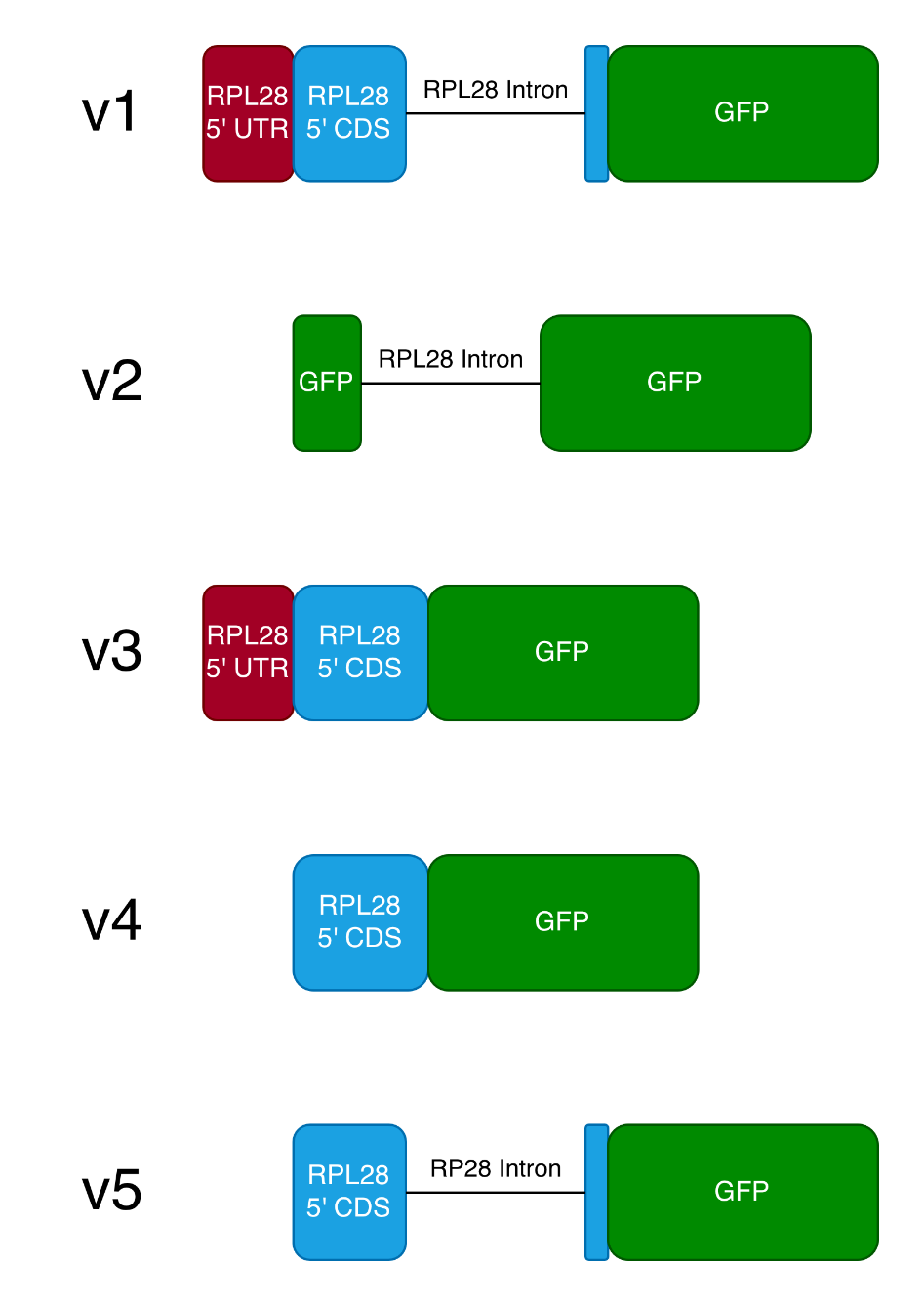

**Supplemental Figure S1. Insertions of RPL28 sequences into GFP.**

V1: original insert containing 16 bases of the 5’ UTR, the first 49 bases of the coding sequence, the intron, and 8 more bases of the coding sequence 5’ to GFP.

V2: Inton only inserted into GFP

V3: same as V1 but without the intron

V4: coding region only, no 5’ UTR or intron

V5: same as V1 without the 5’ UTR

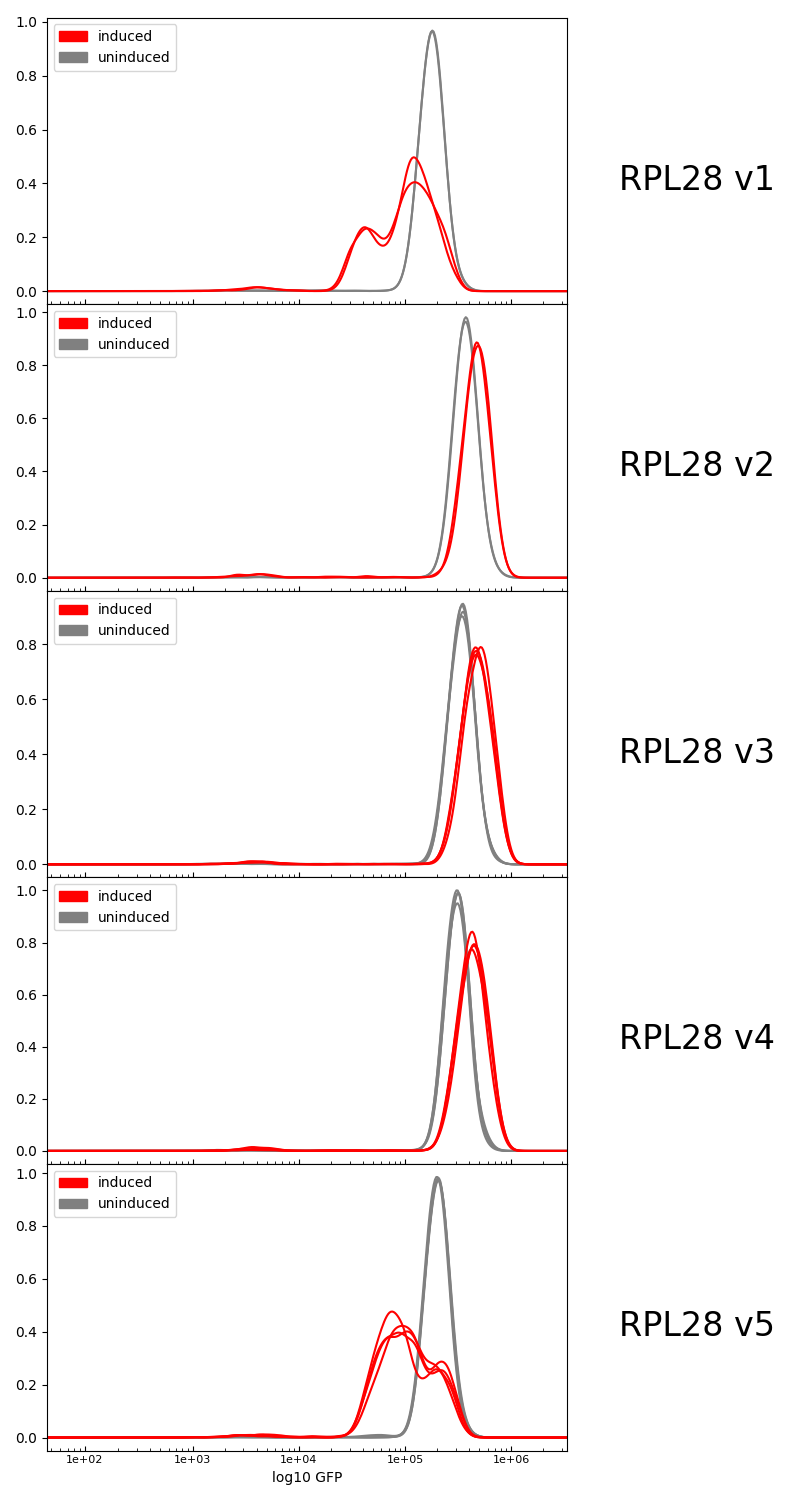

**Supplemental Figure S2: GFP expression for insertion variants**

Horizontal axis is log of GFP fluorescence, vertical axis is number of cells.

The constructs are labeled as in Figure S1.

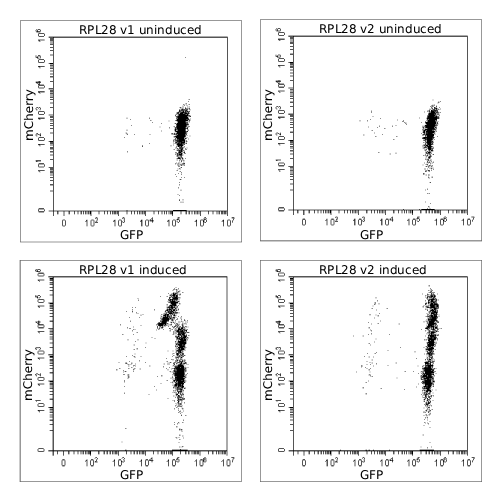

**Supplemental Figure S3: 2D plots of GFP and mCherry fluorescence**

Each dot is a single cell. Plots show both the GFP (horizontal axis) and mCherry (vertical axis) fluorescence values. Left column plots are for construct V1 and right column plots are for construct V2. Upper row in for uninduced condition and lower row is for induced Rpl28 (tagged with mCherry) condition.
